# Thermodynamic analysis of Zα domain-nucleic acid interactions

**DOI:** 10.1101/2022.01.17.476573

**Authors:** Bharath Srinivasan, Krzysztof Kuś, Alekos Athanasiadis

**Author notes:** Both these authors contributed equally to this work (Bharath Srinivasan, Krzysztof Kuś).

## Abstract

DNA/RNA molecules adopting the left-handed conformation (Z-form) have been attributed with immunogenic properties. However, their biological role and importance has been a topic of debate for many years. The discovery of Z-DNA/RNA binding domains (Zα domains) in varied proteins that are involved in the innate immune response, such as the interferon inducible form of the RNA editing enzyme ADAR1 (p150), Z-DNA binding protein 1 (ZBP1), the fish kinase PKZ and the poxvirus inhibitor of interferon response E3L, indicates important roles of Z-DNA/RNA in immunity and self/non-self-discrimination. Such Zα domain-containing proteins recognise left-handed Z-DNA/RNA in a conformation-specific manner. Recent studies have implicated these domains in virus recognition. Given these important emerging roles for the Zα domains, it is pivotal to understand the mechanism of recognition of the Z-DNA/Z-RNA by these domains. To this end, we assessed the binding thermodynamics of Zα domain from ORF112 and ADAR1 on T(CG)_3_ and T(CG)_6_ oligonucleotides which have high propensity to adopt the Z-conformation. Our study highlights important differences in the mode of binding by the two Zα domains originating from different proteins. Site-directed mutagenesis was employed together with isothermal titration calorimetry to tease apart finer details of the binding thermodynamics. Our work advances the understanding on binding thermodynamics of Zα domains to their cognate nucleic acid substrates and contributes to the efforts to gain a complete appreciation of this process.

## Introduction

Nucleic acids are negatively charged biopolymers and their structures are stabilized by base pairing/stacking interactions and metal ion-binding. Nevertheless, nucleic acids show strong conformational fluctuations and are flexible, which is extremely important for their biological functions such as gene replication and expression, protein recognition, and gene regulation (1). The flexibility of nucleic acids strongly depends on their sequence, salt ions in solution, and temperature, which can affect the strength of base pairing/stacking, ion binding, and chain conformational entropy (2). Drugs or proteins can specifically interact with distinct conformations of nucleic acids and can also dramatically affect their structure and flexibility (3, 4). Historically, it has been shown that the double helical structure of DNA adopts three distinct conformations: B-, A- and Z-DNA (5, 6). Z-DNA differs from the other two forms as it adopts the left-handed conformation with 12 base pairs per turn and a steep vertical rise of 3.7 Å per base pair. Additionally, Z-DNA is characterised by the repeating pair of nucleotides with alternating C2’-endo and C2’-exo (compared to a mononucleotide for B- and A-DNA with a sugar pucker that is either C3’-endo or C2’-endo) (5).

It has been shown that Zα domains of several proteins specifically interact with the Z-conformation of nucleic acids (7). It has been demonstrated that Zα domains recognize the characteristic shape of the left-handed helix. This recognition involves interactions with the sugar-phosphate backbone and a conserved CH-π contact formed by a Tyr residue on the protein with a guanosine in syn conformation on the nucleic acid (8–10). Therefore, Zα domains do not discriminate DNA or RNA in left-handed conformation. However, recent results using nuclear magnetic resonance (NMR) has demonstrated that the Zα domains may have kinetic selectivity in binding to either Z-DNA or Z-RNA despite similar binding affinities (11). A complete picture of the macromolecular interaction between the Zα domains and the nucleic acid requires knowledge not only of the structures of co-crystallized complexes of the protein and nucleic acids but also of how the kinetics and thermodynamics drive the binding process. Isothermal titration calorimetry (ITC) studies have been routinely used for studying protein binding to other proteins, peptides, and nucleic acids. Currently, we have partial knowledge of the Zα domain-nucleic acid interaction thermodynamics.

ITC is the method of choice to assess and obtain various thermodynamic parameters such as enthalpy (ΔH), entropy (ΔS), free energy (ΔG), equilibrium binding constant (K_a_), and stoichiometry (12, 13). The binding of a protein to its cognate ligand can be governed by enthalpy, entropy, or both. The enthalpy of binding is driven by non-covalent interactions such as van der Waals interactions (also called London dispersion forces), electrostatic interactions and/or hydrogen bonds. Δ*H* will be negative due to the formation of the energetically favourable noncovalent interactions between atoms of the interacting partners in exothermic reactions. On the other hand, positive enthalpy is due to disruptions of the energetically favourable noncovalent interactions in an endothermic process. However, entropy of binding arises due to reorganization of the backbone or side chain atoms and rearrangement or release of ordered solvent water molecules and ions, resulting in net increase in the entropy of the system (14). The latter is also known as the hydrophobic effect. For a reaction to be thermodynamically favourable, the free energy of the complex formation must be lower than the free energy of interacting partners, i.e., ΔG < 0. Complexes forming due to high entropy changes tend to have weak enthalpy contribution and vice-versa. This phenomenon is called enthalpy-entropy compensation and is common in biomolecular interactions (15). This can be explained by the fact that thermodynamic parameters are intricately interconnected. The high-affinity binding involving noncovalent interactions will lead to a large negative enthalpy change, but it is usually accompanied by a negative entropy difference due to the restriction of the mobility of the interacting partners. Similarly, a large entropy gain is usually associated with an enthalpic penalty (positive enthalpy change) due to the energy required for the disruption of noncovalent interactions (16, 17).

This study attempts to assess the thermodynamics of interaction for the Zα domains of ADAR1 and ORF112 with nucleic acid oligonucleotides T(CG)_3_ and T(CG)_6_. It should be noted that attempts to model the binding of Zα domains to Z-DNA/RNA should consider the B to Z transition (18, 19). Z-DNA binding by Zα domains can be seen as a conformational selection of the Z-DNA form in linked equilibrium (20) (Fig1B) (21). Existence of a left-handed equilibrium and the specific selection model would fit with the idea that antibodies against the Z-form of DNA specifically recognize their cognate epitope rather than bind non-specifically to B-DNA impacting B- to Z-DNA transition. However, there are studies that indicate that Zα domains can bind to oligonucleotide sequences that are in B/A conformation and the binding energy facilitates the B to Z transition provided the oligonucleotide sequences have propensity to adopt Z-conformation (i.e alternating purine-pyrimidine repeats) (22). This study assesses the thermodynamics of binding by Zα domains of ADAR1 and ORF112 and reflects prominent differences in nucleic acid binding by the two proteins. Further, various mutants at the dimer interface and nucleic acid binding pocket of ADAR1 and ORF112 helped tease apart the enthalpic and entropic contributions of protein nucleic acid binding.

**Figure 1.**
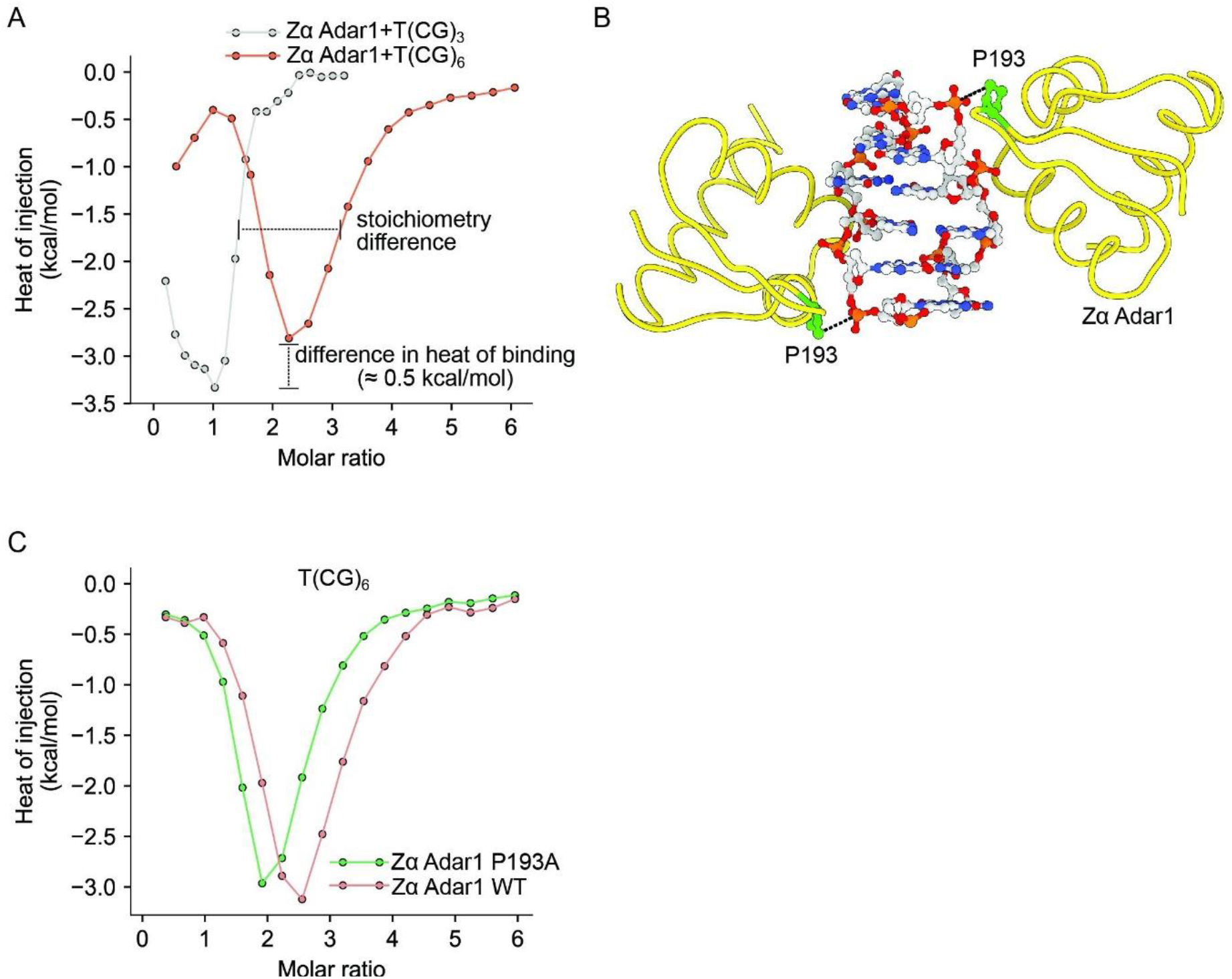
**(A)** ITC profile (Wiseman plot) for Zα ADAR1 binding to T(CG)3 and T(CG)6 oligonucleotide. The filled circles show experimental data points. The lines connect the experimental points and do not represent the model. **(B)** Position of Zα ADAR1 proline 193 residues interacting with the phosphoester backbone of the oligonucleotide. The oligonucleotide is shown in ball and stick representation, the protein is represented as a cartoon **(C)** Superposition of Wiseman plots for Zα ADAR1 wildtype and the P193A mutant binding to the T(CG)6 oligonucleotide.

## Experimental Section and data analysis

### Protein expression and purification

Wildtype proteins and the different mutant constructs were generated, expressed and purified as previously described (8, 23). Briefly, the constructs were transformed into and expressed in *Escherichia coli* BL21 (DE3) strain using kanamycin (50 μg/mL) selection at 37 °C. Cell cultures were induced at mid-log phase (0.6–0.9 OD_600_) with 0.4 mM IPTG. Cells were harvested by centrifugation (5000 × *g*) at 4 °C after 3-4 h post-induction. Cell lysis was performed in buffer containing 20 mM Tris pH 7.6, 50 mM NaCl, 5 mM MgCl_2_, 1X Bugbuster (Novagen) with 1 mM PMSF, proteinase inhibitors cocktail (Complete Mini, EDTA-free; Roche), and benzonase (Novagen) for 1 h at 4 °C and constant stirring. The lysate was centrifuged at 30,597 ×g for 30 min and 0.2 μm syringe filter was used to filter the supernatant. The filtrate was loaded on a HiTrap IMAC-Sepharose FF column (GE Healthcare) pre-equilibrated with buffer A (50 mM Tris pH 7.6, 500 mM NaCl, 1 mM β-mercaptoethanol, 30 mM imidazole). The column was then washed with buffer A, and the protein was eluted using a gradient of 30–500 mM imidazole (buffer B). Overnight dialysis at 4 °C in MonoS buffer A (10 mM HEPES, pH 6.9, 20 mM NaCl) with 10 units of thrombin was carried out to cleave the histidine tag. The resultant protein was loaded on a Mono S 4.6/100 PE (GE Healthcare) pre-equilibrated with MonoS buffer A before washing with a gradient of 20–120 mM NaCl. Protein was eluted with buffer B (10 mM HEPES, pH 6.9, 1M NaCl), and the fractions were evaluated by SDS-PAGE gel electrophoresis. Fractions containing protein were pooled and Amicon-Ultra centrifugal filters (Merck Millipore) were employed for buffer exchange and concentration. The protein was stored in storage buffer (10 mM HEPES, pH 7.4, 20 mM NaCl) at −80 °C and its homogeneity and purity were assessed by SDS-PAGE.

### Isothermal titration calorimetry Experiments

Binding heats were measured on an ITC200 instrument (GE Healthcare) at 25 °C and 1000 rpm. Oligonucleotides T(CG)_3_ and T(CG)_6_ were purchased from Integrated DNA Technologies and annealed. Protein and DNA storage solutions were exchanged against 10 mM HEPES, pH 6.5, 50 mM NaCl with Amicon ultracentrifugal filters (Merck Millipore). Briefly, experiments consisted of 18 injections of 2 μl of protein to oligonucleotide (concentrations used were optimized for optimal curve fit). After each injection, the system was allowed to equilibrate for 3 min.

Raw data were integrated using NITPIC software (24), and fitted with both SEDPHAT (25–27) and/or in CHASM (28). Plots were created with GUSSI (evoked in SEDPHAT) (29) and in CHASM. Figures 1A, 3, 6, S1-S4, S6-S8 is a formal reanalysis of our previously published work (8). Novel data are presented for P193A Zα ADAR1 and S260E, S260Q Zα ORF112.

### Structural analysis

Structure analysis was carried out with Swiss PDB viewer (30) using holo structures for Zα ORF112 (PDB ID: 4wcg) and Zα ADAR1 (PDB ID: 1qbj). For initial assessment, we employed PLIP (31), fully automated protein-ligand profiler, and SCOWLP (32) to extract possible protein ligand interactions for both Zα ADAR1 and Zα ORF 112 for their nucleic acid ligands. Once obtained, the results were critically assessed using detailed analysis performed on PyMol (33).

## Simulation of ITC data

To simulate ITC data that include isomerisation of B- to Z-DNA, we used a method described by Kovrigin (34) that deals with analysis of ITC profiles for systems involving three-state equilibria including both ligand binding and isomerization events. Simulations were performed in MATLAB 2020b using published code with modifications required for newer version of the programming environment. Parameters utilised are included in the supplemental material appendix.

## Results

### Zα ADAR1 Binding to Z-DNA forming oligonucleotides

We were interested in revisiting the binding of Zα ADAR1 to short DNA sequences with propensity to adopt left-handed conformation. To this end, we reanalysed the data presented in Kuś et al.(8) and generated new set of experiments. Examination of the raw calorimetric data reveals that titration of Zα ADAR1 into T(CG)3 during first few injections manifests an endothermic phase followed by an exothermic one (Fig S1). The initial endothermic phase may be indicative of complete or partial desolvation of the duplex due to the introduction of protein that may either mimic salting-out kind of behaviour or of counter-ion stripping mediated by charged groups in protein nucleic acid binding pocket. Further, the heat release reaches plateau, which indicates that all the bound oligo is in the preferred Z-DNA form (Fig 1A). However, the initial endothermic phase cannot be fully explained by the dilution effect (as heat magnitude of protein injection to buffer is at least 3-4 times lower, data not shown). It should be noted that the exothermic and the endothermic phases are not to be confused with spontaneous nature of the reaction since these indicate only changes in Δ*H* (the terms endergonic and exergonic are indicative of changes in Δ*G* which dictates whether reaction is possible or not).

Raw calorimetric data for the titration of Zα ADAR1 into a T(CG)_6_ oligonucleotide containing solution (Fig S2) indicates a biphasic nature with a stoichiometry of 2:1 protein: nucleic acid. Distinct phases may be manifestation of the threshold protein concentration that is required for complete or partial desolvation of the duplex. It should be noted that the concentration equivalence of the lag is 1:1 protein: oligonucleotide and is approximately half of the actual binding stoichiometry. In the light of two binding site, the possibility of a more complex pathway, including an incomplete B-Z transition for the longer oligo, cannot be excluded. Heat exchanged during binding to T(CG)3 is slightly lower than to T(CG)6 (Fig 1A) which might point towards higher number of the favourable bonds created.

Mutations in ADAR1 have been shown to be linked to autoimmune disease Aicardi-Goutières syndrome. One of the distinct *ADAR1* mutations associated with this syndrome is c.577C>G transversion - nonsynonymous Pro193Ala (35). This mutation is localized in the Zα domain. We generated P193A Zα ADAR1 mutant to understand whether changing proline to alanine at the 193^rd^ position causes any substantial difference in the thermodynamics of nucleic acid binding by Zα ADAR1 (Fig 1B). However, only marginal differences were observed between binding of Zα ADAR1 wild-type and the P193A mutant both in terms of stoichiometry and the heat of binding (Fig 1C). This is in agreement with previous study reporting that P193A had only a minor reduction in binding affinity compared to WT (36).

### Different mode of Zα ORF112 Binding to Z-DNA forming oligonucleotides

Titration of Zα ORF112 into a T(CG)_3_ oligonucleotide containing solution shows a clear exothermic transition (Fig S3). Binding of Zα ORF112 to the T(CG)_3_ oligo is more exothermic and indicates a sharper transition than the one observed for Zα ADAR1 to T(CG)_3_ (Fig 2A). It would be expected that titrating Zα ORF112 into a solution containing the T(CG)_3_ oligonucleotide should be also enthalpically unfavourable due to possible desolvation. However, this might be compensated by the enthalpically favourable binding of the protein to the DNA backbone. Thus, the absence of the initial endothermic phase could be explained by enthalpy-entropy compensation. Given the higher enthalpy of binding of the oligonucleotides for Zα ORF112, the entropy is restricted due to possible molecular constraints that results in compensatory conformational entropy reduction. This phenomenon is well documented in literature for several different protein-ligand interactions (16). Titration of Zα ORF112 into a T(CG)_6_ exhibits the same behaviour as seen with Zα ORF112 and T(CG)_3_. However, the stoichiometry between protein: nucleic acid is doubled indicating two binding sites on the longer oligonucleotide (Fig 2B).

**Figure 2.**
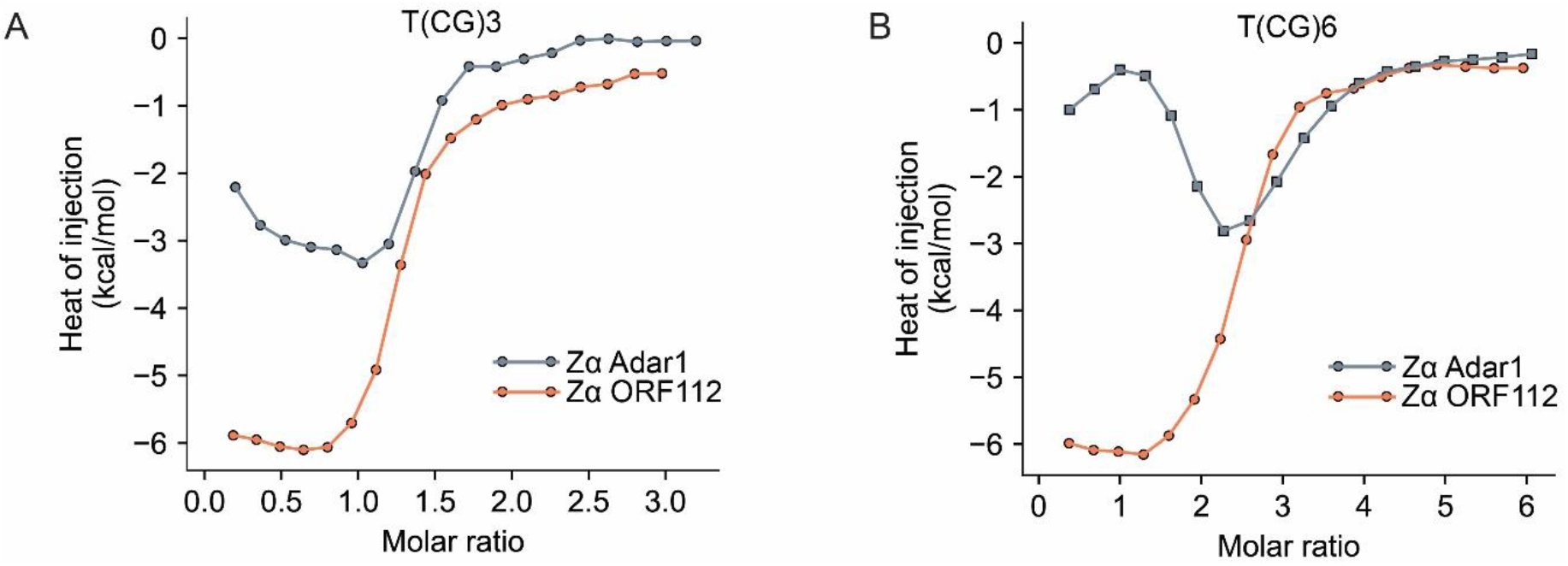
**(A)** Superposition of Wiseman plots for Zα ADAR1 and Zα ORF 112 binding to T(CG)_3_ and **(B)** T(CG)_6_.

### Potential explanations of differential nucleic acid binding by Zα ADAR1 and ORF 112

As mentioned, Zα ADAR1 and Zα ORF 112 binding to both T(CG)_3_ and T(CG)_6_ oligonucleotide display differences (Fig 2A, B). The magnitude of enthalpy change in the exothermic phases for T(CG)_3_ and T(CG)_6_ is almost 3-fold higher for the Zα ORF112 compared to Zα ADAR1. However, the total ΔG might remain relatively unchanged given that the reduced ΔH is compensated by the gain in entropy resulting in comparable affinities (Table S1). However, the shapes of the ITC plots suggest dissimilar entropic and enthalpic contributions that may reflect differences in the binding mechanisms.

Sequence alignment of Zα domains of ADAR1 and ORF 112, shows ~28 % sequence conservation including residues that are directly implicated in nucleic acid binding (Fig 3A). However, there are two plausible explanations for such pronounced differences in the ITC profiles between the binding of oligonucleotide by Zα ADAR1 and Zα ORF 112:

**Figure 3.**
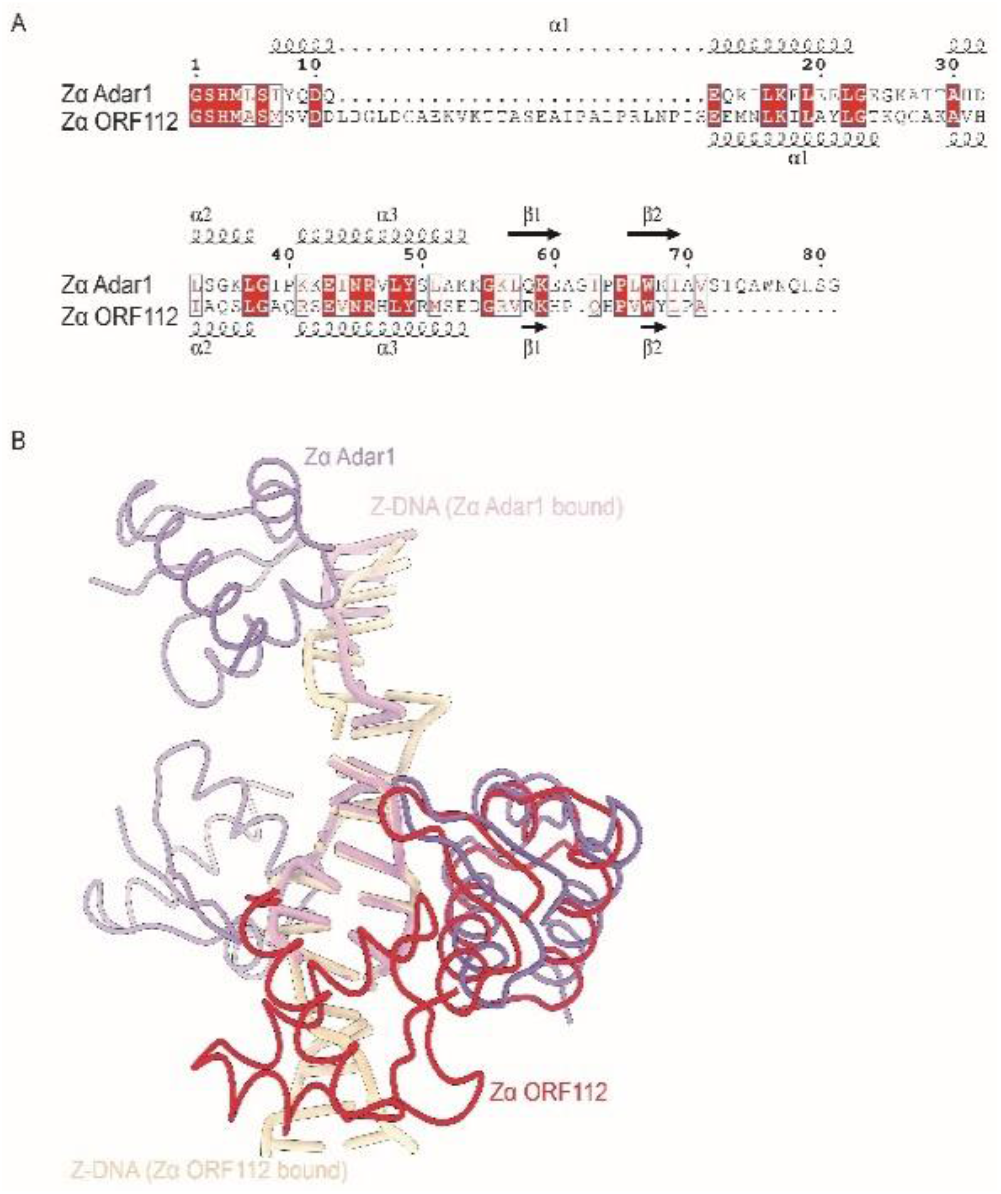
**(A)** Sequence alignment of Zα ORF112 (PDB ID: WCG) and Zα ADAR1(PDB ID: 1QBJ). Alignment was generated in MUSCLE and rendered in ESPrint. **(B)** Superposition of structures for Zα ORF112 (PDB ID: 4WCG) (red)) and Zα ADAR1 (PDB ID: 1qbj) (violet).

A) The different mode of binding. Superimposition of the structures of Zα ADAR1 and Zα ORF 112 shows Zα ADAR1 binds to the oligonucleotide by sandwiching the DNA on both the strands. However, Zα ORF112 seems to be binding to the DNA at a single site that propagates longitudinally along the axis of the nucleic acid on one side by means of protein oligomerization that is possibly mediated by nucleic acid (Fig 3B).

B) Differences in the interactions and interface water-mediated hydrogen bonds between Zα ADAR1 and Zα ORF 112. Since we observe an initial endothermic phase in the curves of Zα from ADAR1, we were particularly interested in interactions that could possibly indicate differences in either desolvation or ion-stripping effect. To this end, we used PLIP, fully automated protein-ligand profiler, and SCOWL (see methods) to extract possible protein ligand interactions for both Zα ADAR1 and Zα ORF 112. Our analysis indicates substantial differences between Zα ADAR1 and nucleic acid interaction compared to Zα ORF112. It should be noted that the interactions discussed here are for the C chain of protein with the F-chain of nucleic acid for 1QBJ (ADAR1); B chain of protein with C chain of nucleic acid for 4WCG (ORF112). It should be noted that the A chain in 4WCG (ORF112) makes qualitatively different contacts with the nucleic acid which are not presented here.

Zα domains differs in the network of hydrogen bonds formed with the nucleic acid. Hydrogen bond is an electrostatic attraction between two polar groups that occurs when a hydrogen atom is shared between two electronegative atoms. Hydrogen bond strength will depend on distance between the donor and the acceptor atoms and the angle between them. Zα ADAR1 utilises residues K169, N173, R174 and T191 to create hydrogen bonds with the nucleic acid, whereas Zα ORF112 uses S250, N253 and R258 to form these interactions. Water forms bridging networks and plays critical role in dictating the binding mode of ligands and is emerging as an important determinant of binding thermodynamics. A water bridge is defined as a contact between two heavy atoms (nitrogen or oxygen) of the macromolecules mediated by one water molecule through two hydrogen bonds. K169 and K170 of Zα ADAR1 make important water bridges with the nucleic acid ligand (the equivalent residues in the B chain of protein in Zα ORF112 are R249 and S250, both of which are not involved in many water-mediated interactions). However, the only water bridge mediated interaction detected for Zα ORF112 is through Y257 which is also involved in π-stacking (the equivalent residue in Zα ADAR1, Y177, also makes π-stacking interaction with nucleic acid but fails to make a water bridge interaction).

Salt bridges are non-covalent interactions between two ionized functional group with a hydrogen bond and an electrostatic interaction component. Zα ADAR1 uses K169, K170 and R174 to create salt-bridge interaction with the phosphate groups. However, the only group that is involved in salt-bridge interaction in Zα ORF112 is R254. Overall, these thermodynamic and structural analyses indicate that interaction modalities of Zα domains from ADAR1 and ORF112 are substantially different.

As mentioned above, DNA-bound Zα ORF112 monomers make extensive contacts through their α3 helix and the wing region. To understand the differential binding displayed by Zα ADAR1 and Zα ORF 112 and to tease apart the role of protein oligomerization in DNA binding, a series of mutants were reanalysed or created. Serine 260 is a residue very close to the protein-protein interface of the on-DNA oligomerized Zα ORF112 (Fig 4A). Perturbation of this interaction should shed light on the possible role of protein oligomerization in DNA binding. With that aim, serine 260 was mutated to glutamate residue to create unfavourable electrostatic barrier for oligomerization. Since this residue is far from the nucleic acid interaction site, it is unlikely that it will interfere strongly with protein-nucleic acid interaction. Further, since this residue is flanked by glutamate 261 and aspartate 262, mutating S260 to glutamate will concentrate too much negative charge on the interface to offset any nucleic acid mediated interaction. Inspection of curves in Fig 4B and S5 indicates that on-DNA dimerization is an essential aspect for high affinity binding by Zα ORF112. Breaking the dimerization with accumulation of negative charge causes the ITC titration curve to look like that displayed by Zα ADAR1 (Figure 5A, B, C, D).

**Figure 4.**
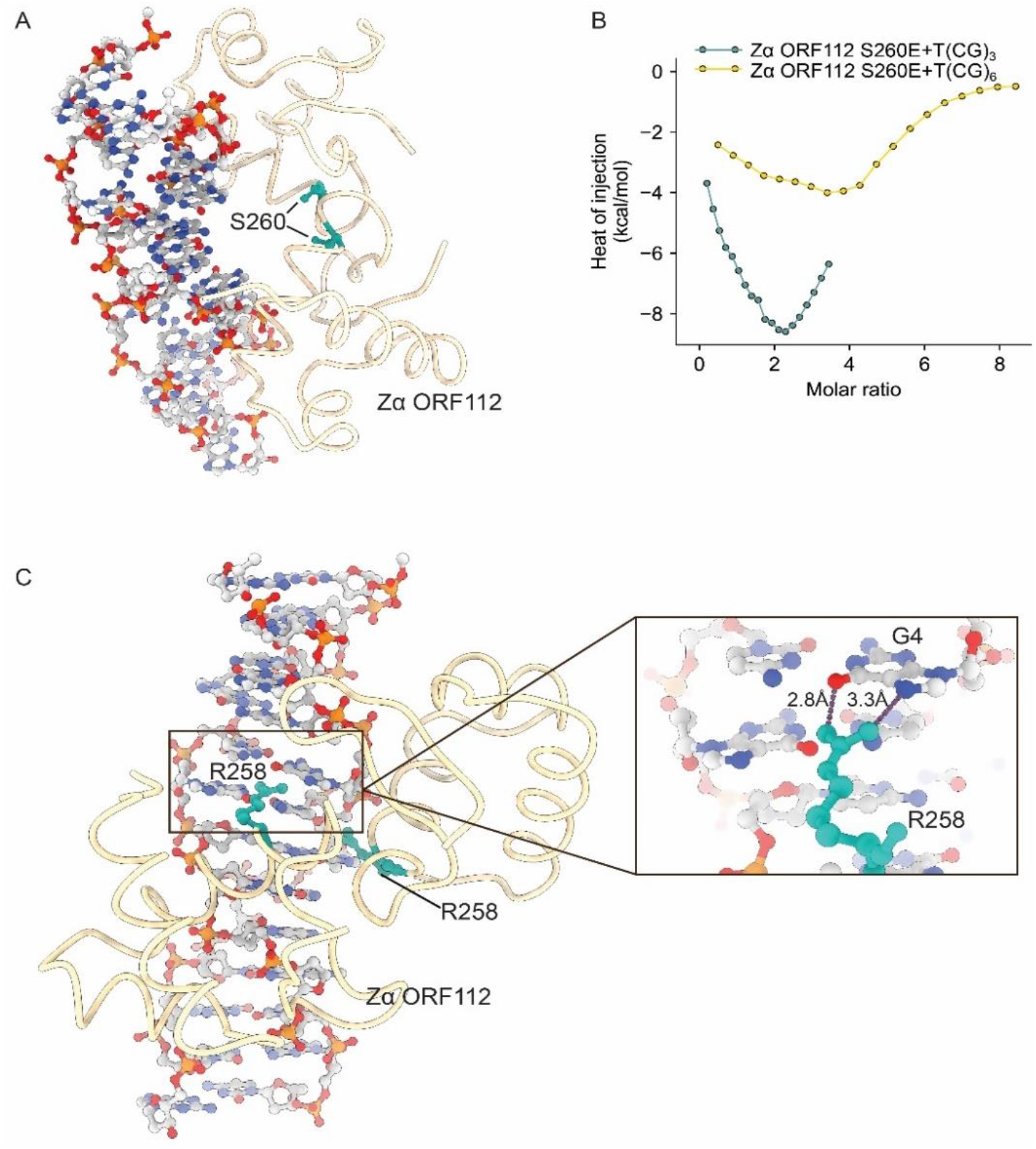
**(A)** Zα ORF112 on-DNA dimer. The serine 260 residue at the protein-protein interface is depicted as green ball and sticks. **(B)** Integrated heats of binding (Wiseman plots) for T(CG)_3_ and T(CG)_6_ binding by ORF112 S260E. **(C)** Representation of the interaction between Zα ORF112 arginine 258 and nucleic acid. Right panel shows the zoom in of R258 residue contacting the C6 carbon of the guanine residue (G4).

**Figure 5.**
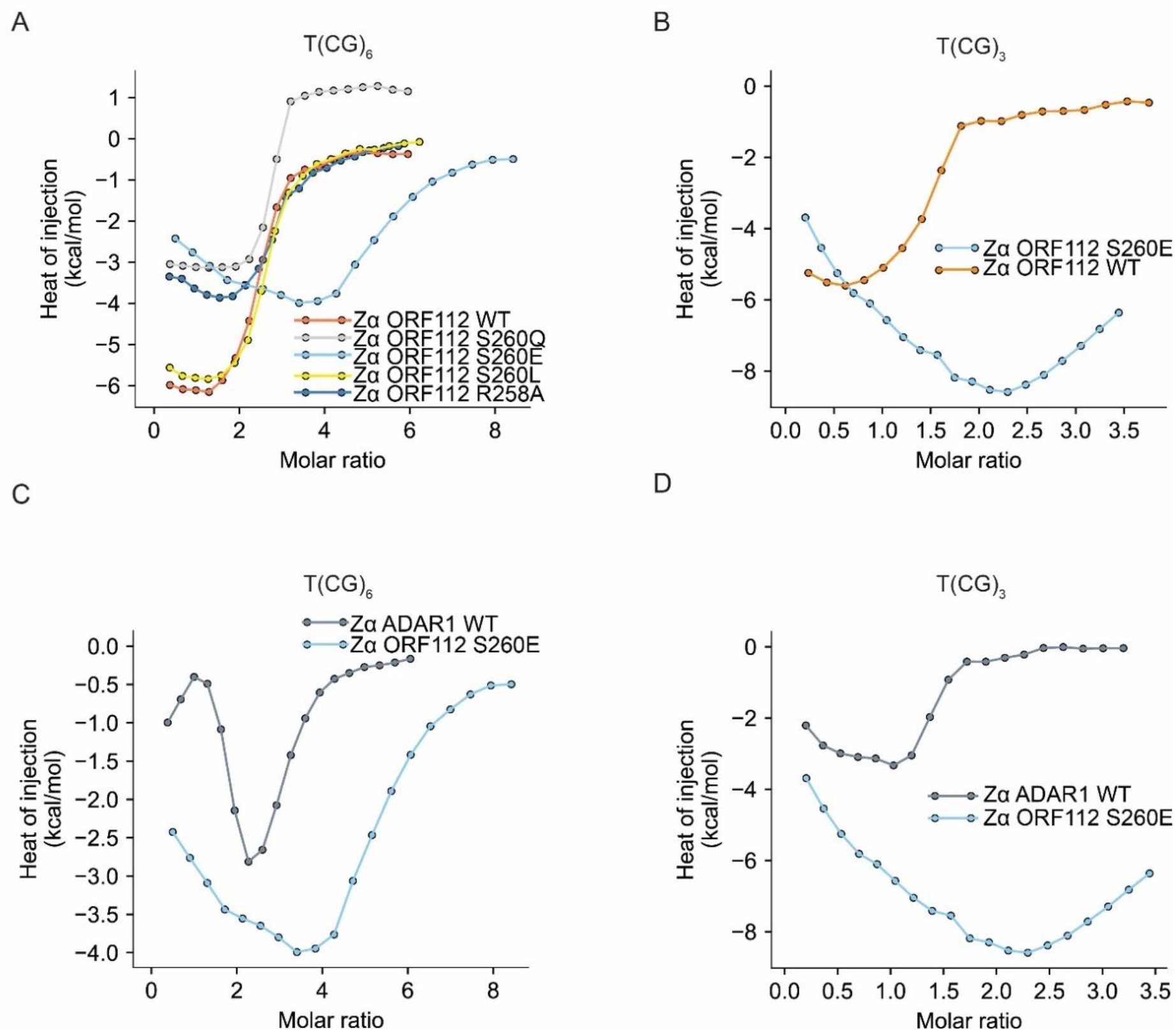
**(A)** Superposition of Wiseman plots for Zα ORF112 wild-type and its mutants (S260E, S260L, S260Q and R258A) to T(CG)_6_. **(B)** Superposition of ITC curves for wild-type and the mutant variant S260E of ORF112 binding to T(CG)_3_ **(C)** Superposition of ITC curves for wild-type Zα ADAR1 and the mutant variant S260E of Zα ORF112 binding to T(CG)_6_. **(D)** Superposition of ITC curves for wild-type Zα ADAR1 and the mutant variant S260E of ORF112 binding to T(CG)_3_.

However, the affinities are poorer and the stoichiometry of protein: nucleic acid is larger than that observed for Zα ADAR1. For Zα ORF112 S260E, the binding stoichiometry is 4:1 protein: nucleic acid for the T(CG)_6_ oligonucleotide and 2:1 for T(CG)_3_ oligonucleotide (Fig 5A, B, C and D). This could be possibly explained by the fact that ORF 112 is a constitutive dimer and has two sites of binding for the dimer on the oligonucleotide. This will essentially result in 4 sites of binding for the monomer once the dimer dissociating mutant is created. This observation could be substantiated by high ionic strength induced dimerization seen in the presence of ammonium sulphate, a mimic of phosphate. The oligonucleotide containing solution, into which the protein is injected, might mimic ammonium sulphate and result in high ionic strength inducing dimerization. Further, it should be considered that ITC experiments are carried out at an order-of-magnitude higher concentrations compared to gel filtration experiments (the protein gets rapidly diluted in the mobile phase of a gel filtration experiment). Oligomerization, to a large extent, is a property of the protein concentration especially in scenarios where there is a dimer-monomer equilibrium.

To validate that the observed behaviour for Zα ORF112 S260E is exclusively charge dependent and is not a property of the size of the substituted amino acid, we mutated serine 260 to glutamine (the amide group of glutamic acid is neutral under all biological conditions). Further, it is a polar residue similarly to the serine. ITC profiles for the binding of Zα ORF112 S260Q into a solution containing the T(CG)6 oligonucleotide (Fig 5A, S7) shows that the overall shape of the normalized curve resembles that of the wild-type ORF112. However, careful inspection of the power consumption for maintaining the baseline seems to be slightly endothermic at saturating proteins indicating competing entropic contributions. It should be noted that though glutamine is neutral, it has a long R group possibly causing steric hindrance locally that has overall impact on hydrogen bonding residue alignment.

On the other hand, mutating the serine to leucine might promote a tighter dimer due to solvent exclusion and hydrophobic interaction, thus promoting protein nucleic acid interaction similar to or better than the wild type. These predictions are in line with the revisited data for Zα ORF112 S260L indicating that its affinity is comparable or slightly better than WT (Fig 5A, S6).

To understand whether perturbation of base specific contacts of the protein with nucleic acid will modulate the binding behaviour, we reanalysed the data for Zα ORF112 R258A. Arginine 258 is within hydrogen bonding distance (~2.8 Å) of the C-6 carbonyl oxygen of guanine nucleobase (Fig 4C). Mutation of the residue would perturb the hydrogen bonding and will reflect the importance of such specific enthalpically driven interactions on the overall thermogram of binding. In fact, the overall shape of the thermogram does not change (Fig 5A, S8). However, the ΔH changes and becomes less steep indicating poorer binding than the wild type.

### Modelling of the isothermal titration calorimetry data that includes B-to-Z transition

Although there is evidence showing that Zα domains can bind to both left-handed and right-handed conformation of nucleic acids, for simplicity, we consider only binding to Z-conformation (all-or-none Monod-Wyman-Changeux model, MWC) (Fig 6A). Since an induced fit sequential model like KNF (Koshland, Némethy and Filmer model, KNF model) or complex pathways (Fig 6B) require more elaborate mathematical treatment, we opted to use the MWC model to approximate the binding behaviour. Further, the MWC model has a smaller parameter space which has lower risk of over-fitting.

**Figure 6.**
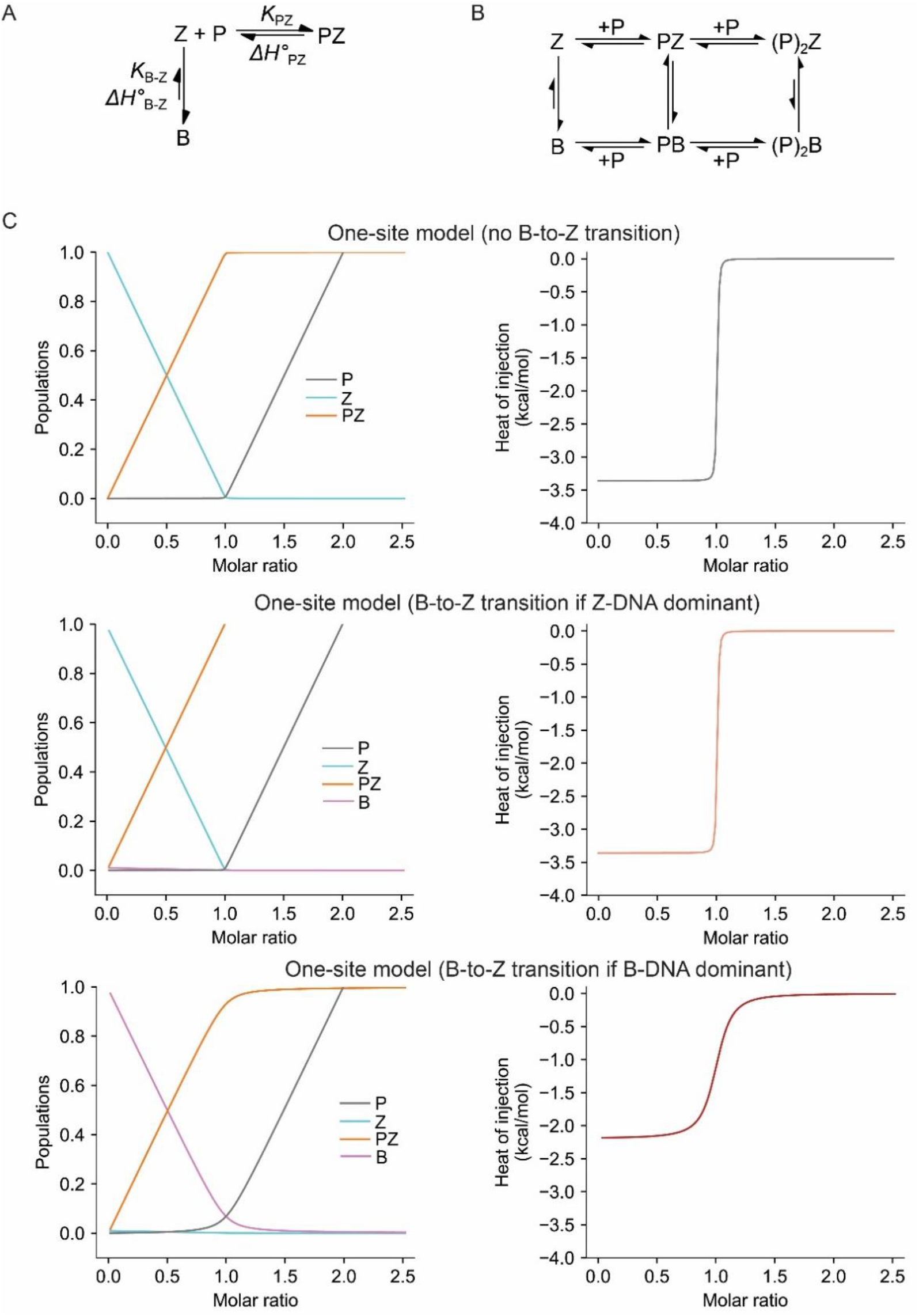
**(A)** Model describing the binding of Z-DNA by Zα domains including conformational changes of nucleic acid where P is the Zα domain (protein), Z – the Z-DNA ligand, B – B-DNA, and PZ is the macromolecule-ligand complex. K_PZ_ and ΔH°_PZ_ are the intrinsic energetics for ligand-macromolecule binding. K_B-Z_ and ΔH°_B-Z_ are parameters describing conformational switching between the B-form and the Z-form of the oligonucleotide. **(B)** Schematic representation of the potential pathways (considering both KNF- and MWC-models of induced fit and population shift type transitions) through which Z-DNA/RNA-protein complex can be formed. Abbreviations used as in (A). **(C)** Simulations of isothermal calorimetry data assuming variants of the model from panel (A). Left panel shows the species changes and right one the heat of injections (although set as kcal/mol, this units are arbitrary). Parameter used are included in supplementary material as Appendix.

Analysis of calorimetric data rely on the fact that equilibrium constants *(K)* and enthalpy changes (Δ*H*) are observed equilibrium constants, *K*_obs_, and observed enthalpy changes, Δ*H*_obs_, as they include contributions from a series of binding related phenomena. These binding events range from the assumed trivial ones including removal of solvent to more complex coupled events like release of counterions or (de)protonation. Since, in the current model nucleic acid conformational change and binding are linked (Fig 6A), the observed binding constant, *K*_obs_, includes contributions from both the binding and conformation transition equilibria. It is defined as follows:

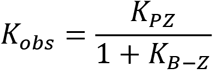

where *K_PZ_* is the intrinsic binding constant for binding the ligand to the Z conformation of the nucleic acid molecule and *K*_B-Z_ is the transition equilibrium constant for the nucleic acid to go from B to Z conformation. The observed enthalpy is described by:

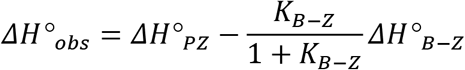

Where ΔH°_PZ_ is the intrinsic enthalpy for ligand-macromolecule binding, the ratio *K*_B-Z_/ (1+ *K*_B-Z_) describes the fraction of nucleic acid in the B-conformation, and ΔH°_B-Z_ is the enthalpy of B to Z transition.

To explore how inclusion of the B-to-Z transition might impact the profiles of the ITC, we utilised a theoretical approach developed by (34) which uses differential equations to simulate changes in the concentrations of the different species during the injections, which in turn allows (given enthalpy) to generate Wiseman plot. In first instance, we simulated one site binding model (excluding B-to-Z) (Fig 6C, top panels). Left side of the figure presents the changes in the populations of the species and right one illustrates the heat changes during titration. To test how inclusion of the B-to-Z transition affects the predicted enthalpy changes, we included B-to-Z transition and considered a scenario when Z-DNA is dominant (Fig 6C, middle panel). The behaviour, as expected, is indistinguishable from the model that does not include B-to-Z transition. Lastly, we simulated the model when B-DNA is dominant form (Fig 6C, bottom panel) which changes the profiles of the heats. Nevertheless, it should be noted that curve shape has resemblance of the simple one site-model which has been described in the theoretical analysis (34).

## Discussion

Modelling the interaction of Zα domains with DNA and RNA oligonucleotides poses multiple challenges. One of the issues is knowledge of the exact concentration of the Z-DNA. In low salt and room temperature, the equilibrium is shifted towards the B form. This is even more important given the continuing debate about the mode of Zα domain binding to DNA/RNA. Some studies support the notion of induced fit (KNF model) (37) where Zα domains bind to the B form of DNA and induce the B- to Z-transition that is subsequently stabilised by on DNA dimerization of the protein (Figure 6B). Other studies favour the model of Zα domains exclusively recognizing the Z-conformation of nucleic acid to bind and shift the equilibrium in a time-dependent manner (Monod-Wyman-Changeux model, MWC model). It has been demonstrated that (dCdG)_4_ remains in B-form (detectable population in the Boltzmann distribution is B-form) at 25 °C in 10 mM Phosphate buffer both at 2.0 M NaCl or at 115 mM Na^+^ and 200 μM [Co(NH_3_)_6_]^3+^. Hence it is highly likely that only a very minor fraction (<10%) of T(CG)_3_ and T(CG)_6_, the nucleic acid oligonucleotides employed in the present study, can adopt Z-DNA/RNA conformation at 25 °C in 10 mM HEPES, pH 6.5, 50 mM NaCl. This minor Z-form in the population gets captured and trapped thus, shifting the equilibrium towards all Z-DNA by approximately 30 minutes of protein titration. Previous studies have reported a *ΔH°* of ~700 cal/mol for the (dCdG)_4_ oligonucleotide under conditions of increasing monovalent or polyvalent salt (38). Further, another study using single-molecule Fluorescence Resonance Energy Transfer (FRET) assays for Z-conformation adopting sequences embedded in a long inactive DNA, measured the thermodynamic populations distribution of ADAR1-bound DNA conformations in both GC and TG repeat sequences. Based on a statistical physics model, they predicted the affinities of ADAR1 to both Z-form and B-form of these sequences and suggested the presence of an intermediate B* state going from B-Z conformation (22).

Another potential problem in modelling the binding is the information rich nature of the binding isotherms. As seen in the curves, Zα ADAR1 binding to DNA oligonucleotides has an initial endothermic phase followed by an exothermic phase. The endothermic phase becomes more pronounced with increasing length of the oligonucleotide (T(CG)_6_ > T(CG)_3_). The initial endothermic phase (with negative Δ*H*), indicative of possible desolvation or charged counterion stripping from the charged phosphate of the nucleic acid, cannot be explained by current models (39, 40). Breaking of bonds, even solvent, is endothermic. Solvent distribution would have to be modelled to understand the endothermic phase. This is important given the clear role of entropy as the driving force in binding of protein onto nucleic acids in the case of monomeric variants of ORF112 and ADAR1 versus dimeric WT ORF112. Analysis of water network in Zα ADAR1 and ORF112 structures at the interface with their respective oligonucleotide ligands indicates that the water distribution is not conserved possibly leading to differential mode of binding as is evident in the ITC data. Additionally, dimer-monomer equilibrium usually complicates the modelling of ITC data (41). In our case, we know that ORF112 possibly interacts with the nucleic acid as a dimer (8). This manifests itself when dimer destabilizing mutant doubles the stoichiometries in the ITC data. Thus, incorporating the association and dissociation rates (and hence, the rate constants) of the dimer in modelling the thermodynamics of protein nucleic acid interaction is very important.

It has been repeatedly discussed in literature that antibodies against Z-DNA/RNA detects the nucleic acid only downstream of RNA polymerase moving site (42), possibly indicative of some role for protein in Z-DNA formation and maintenance. This gives rise to the question whether Zα domain containing proteins induce the Z conformation in oligonucleotides with the propensity to form such structures? Though, we do not have conclusive evidence either to support or refute this possibility, it will run contrary to the understanding that Zα domain containing proteins specifically localize to the Z-DNA containing nucleotide stretches. Further, we also know that antibodies against Z-DNA specifically recognize the Z-conformation rather than brining about B-Z transition. Hence, the premise of the present study is the assumption that Zα domain containing proteins bind to Z-DNA/RNA and this assumption remains invariable until further experimental studies provide conclusive evidence on protein induced Z-DNA/RNA formation.

Modelling of the ITC data is non-trivial but from theoretical point of view inclusion of the isomerisation step scales the concentrations of components during the titration but resulting theoretical Wiseman plots still allows to fit the one site model (34). Therefore, models with linked equilibrium might not be easily spotted in the ITC analysis. Nevertheless, to produce sigmoidal shaped Wiseman plot it is necessary that K_B-Z_ ≪ K_PZ_. Therefore, to fully understand binding of Z-DNA by Zα domain integrative approach will be required which includes multiple experimental setups. ITC, although powerful technique providing multiple thermodynamic parameters, might be insufficient to distinguish finer modes of binding if the model can be described by one-site binding. In conclusion, this study represents an exploration of different binding modes by Zα domains and is an outlook at modelling of the isothermal titration data in the context of Z-DNA binding.

## Abbreviations

ITC: Isothermal titration calorimetry
ADAR1: Adenosine deaminase acting on RNA 1
ZBP1: Z-DNA binding protein 1
MWC model: Monod-Wyman-Changeux model
KNF model: Koshland-Nemethy-Filmer model

## Dedication

This manuscript is dedicated to the memory of Alekos Athanasiadis who sadly passed away in August 2020. Though the manuscript is communicated by BS and KK for practical purposes, the work was conceived and supervised by AA.

## Acknowledgements

We thank Élio Sucena, Jonathan C. Howard and Mónica Bettencourt-Dias for providing useful comments. This work was supported by a Fundação para a Ciência e Tecnologia (FCT) grant PTDC/BBB-BEP/3380/2014 to BS and AA. BS was also supported by Marie Skłodowska-Curie Individual Fellowship Grant 789565. KK was a recipient of FCT PhD grant SFRH/BD/51626/2011.

## Notes

### Competing Interest Statement

The authors have declared no competing interest.

