## Supplemental material for "Thermodynamic analysis of Zα domain-nucleic acid interactions"

Current address, [c]Mechanistic Biology and Profiling, Discovery Sciences, R&D, AstraZeneca, Cambridge, UK; [d]Department of Biochemistry, University of Oxford, South Parks Road, Oxford, OX1 3QU, UK

[b]Corresponding authors;

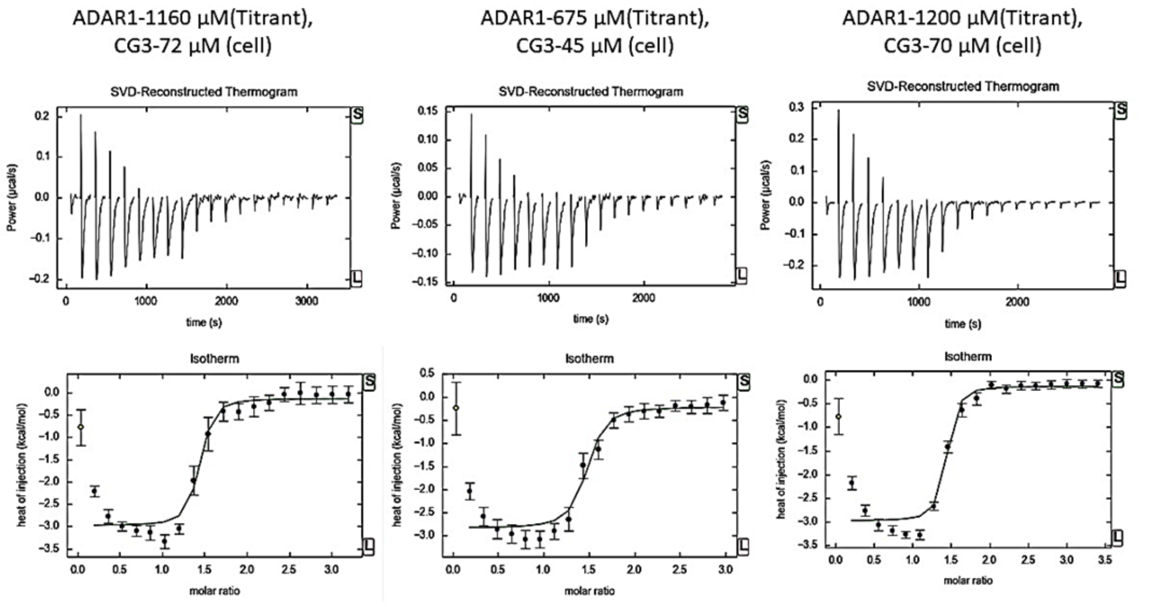


**Figure S1.** Isothermal titration calorimetry profiles for the titration of Zα ADAR1 into a solution containing the $T(CG)_3$ oligonucleotide at various ratiometric concentrations of the protein and the oligonucleotide. The raw calorimetric titration data for the injections (2μl/injection) are shown in the upper panel and the integrated results after subtraction of heats of dilution (Wiseman plots) are shown in the lower panels (as circles) and the solid lines represent the best least-square fits for the integrated data. The data in the above panels are mostly fitted to one site binding ignoring the endothermic phase completely. For column 1, approximate equimolarity between the nucleic acid and the protein would be achieved after 10 injections, for column 2 it will be after 11 injections, and for column 3 it will be after 10 injections (one titration takes approximately 54 minutes).

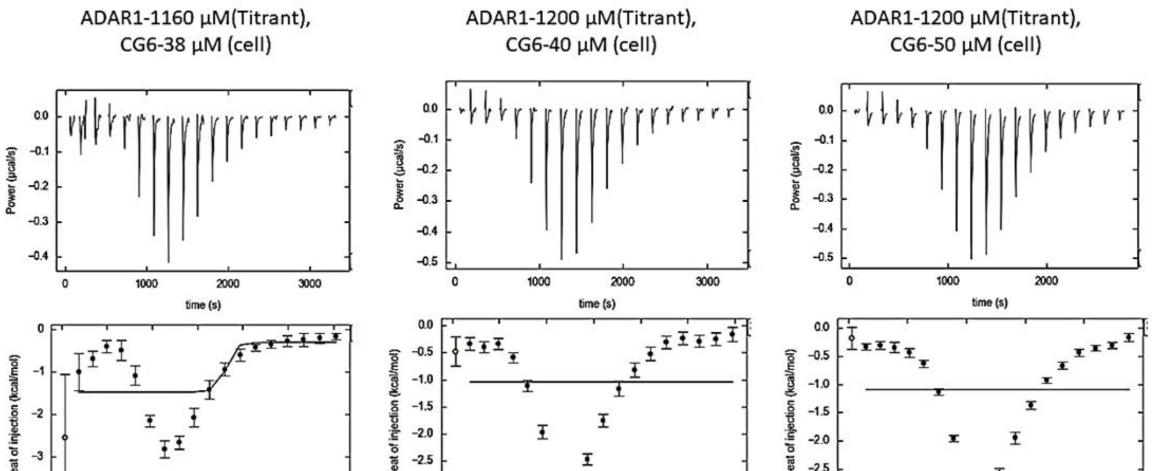


**Figure S2.** Isothermal titration calorimetry profiles for the titration of Zα ADAR1 into a solution containing the $T(CG)_6$ oligonucleotide at various ratiometric concentrations of the protein and the oligonucleotide. The raw calorimetric titration data for the injections (2μl/injection) are shown in the upper panel and the integrated results after subtraction of heats of dilution (Wiseman plots) are shown in the lower panels (as circles) and the solid lines represent the best least-square fits for the integrated data. The data in the above panels are mostly fitted to one site binding ignoring the endothermic phase completely.

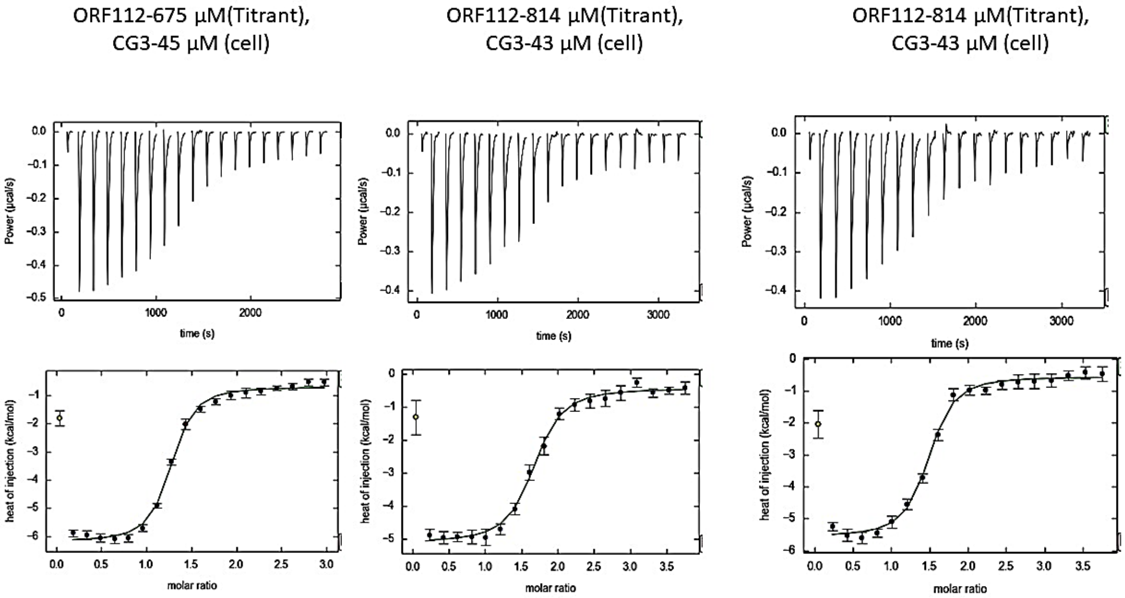


**Figure S3.** Isothermal titration calorimetry profiles for the titration of ORF112 into a solution containing the $T(CG)_3$ oligonucleotide at various ratiometric concentrations of the protein and the oligonucleotide. The raw calorimetric titration data for the injections (2µl/injection) are shown in the upper panel and the integrated results after subtraction of heats of dilution are shown in the lower panels (as circles) and the solid lines represent the best least-square fits for the integrated data. The data in the above panels are mostly fitted to one site binding ignoring the endothermic phase completely.

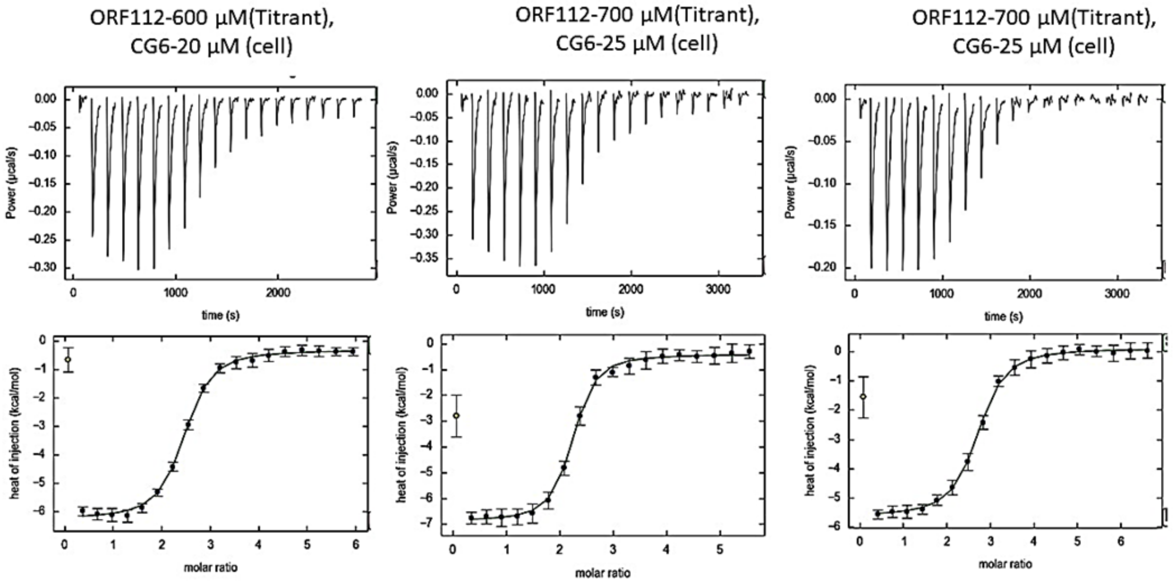


**Figure S4.** Isothermal titration calorimetry profiles for the titration of ORF112 into a solution containing the $T(CG)_6$ oligonucleotide at various ratiometric concentrations of the protein and the oligonucleotide. The raw calorimetric titration data for the injections (2µl/injection) are shown in the upper panel and the integrated results after subtraction of heats of dilution are shown in the lower panels (as circles) and the solid lines represent the best least-square fits for the integrated data. The data in the above panels are mostly fitted to one site binding ignoring the endothermic phase completely.

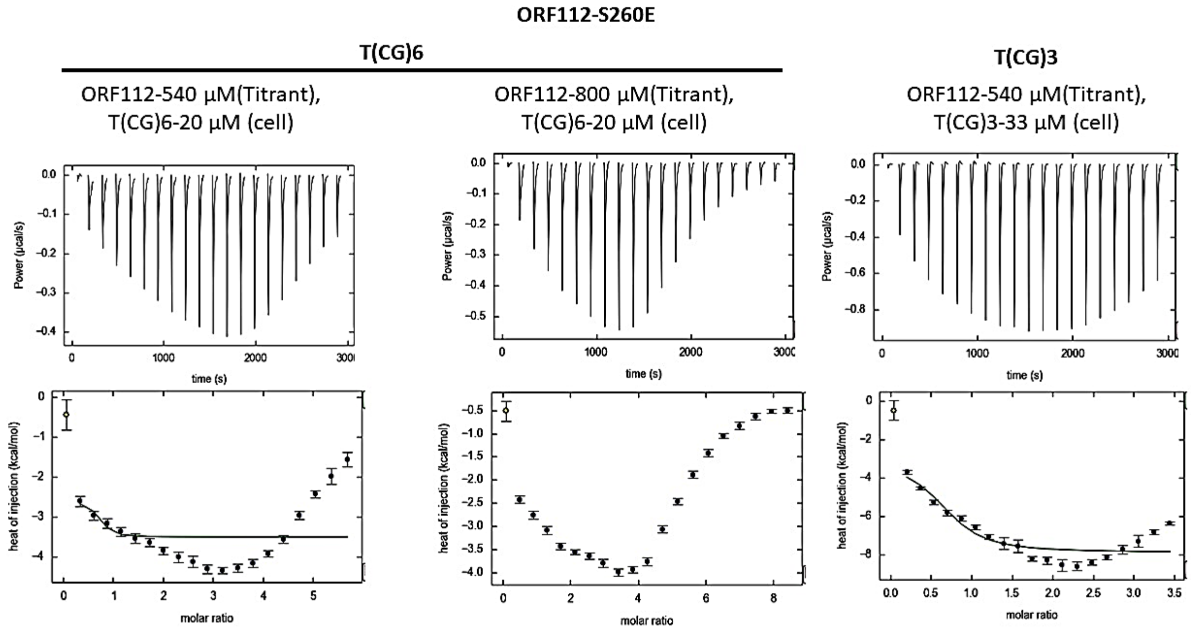


**Figure S5.** Isothermal titration calorimetry profiles for the titration of Zα ORF112 S260E into a solution containing the T(CG)$_6$ oligonucleotide and T(CG)$_3$ oligonucleotide at various ratiometric concentrations of the protein and the oligonucleotide, respectively. The raw calorimetric titration data for the injections (2μl/injection) are shown in the upper panel and the integrated results after subtraction of heats of dilution are shown in the lower panels (as circles) and the solid lines represent the best least-square fits for the integrated data.

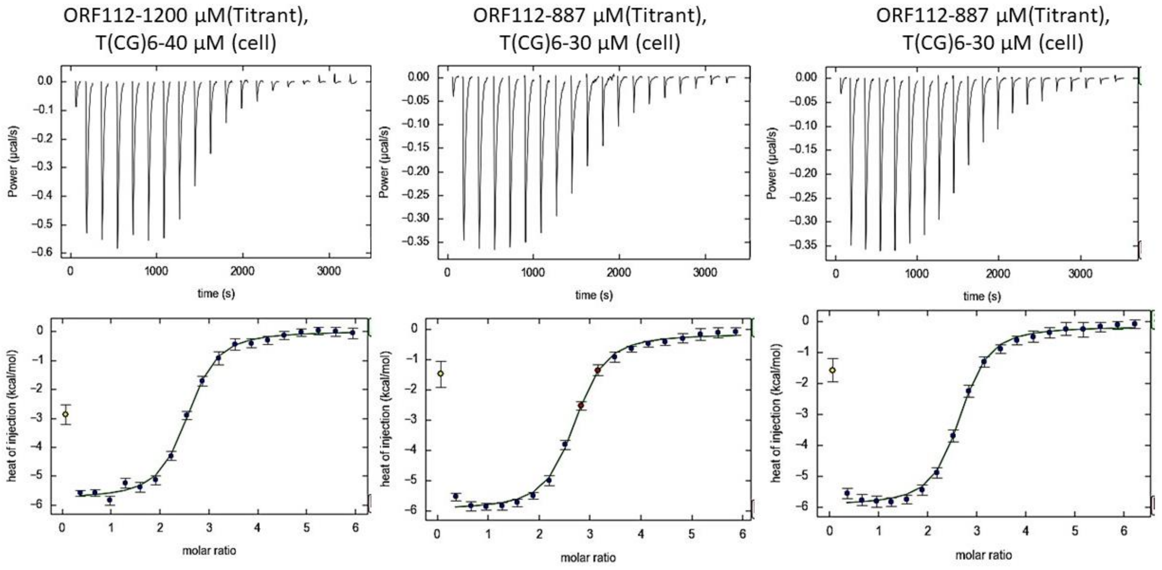


**Figure S6.** Isothermal titration calorimetry profiles for the titration of Zα ORF112 S260L into a solution containing the T(CG)$_6$ oligonucleotide at various ratiometric concentrations of the protein and the oligonucleotide, respectively. The raw calorimetric titration data for the injections (2μl/injection) are shown in the upper panel and the integrated results after subtraction of heats of dilution are shown in the lower panels (as circles) and the solid lines represent the best least-square fits for the integrated data.

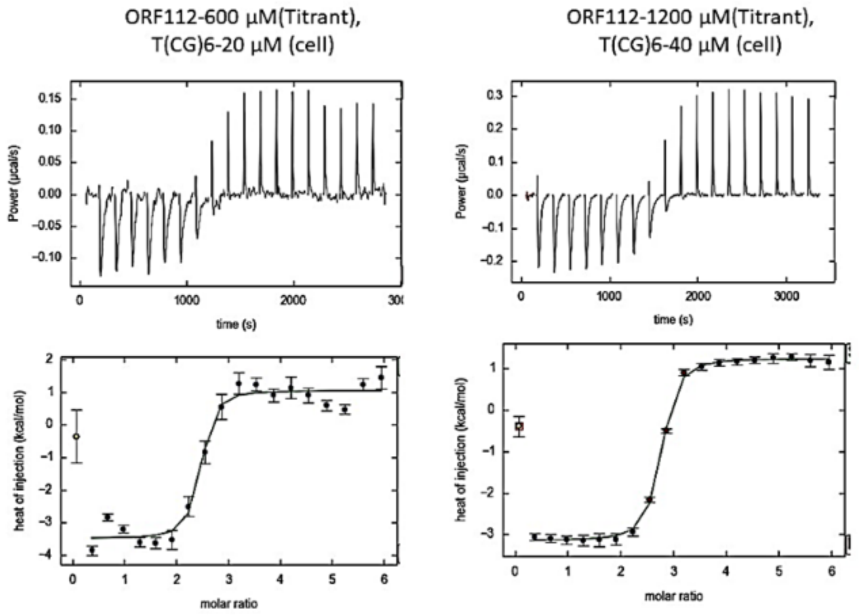


**Figure S7.** Isothermal titration calorimetry profiles for the titration of Zα ORF112 S260Q into a solution containing the $T(CG)_6$ oligonucleotide at various ratiometric concentrations of the protein and the oligonucleotide, respectively. The raw calorimetric titration data for the injections (2µl/injection) are shown in the upper panel and the integrated results after subtraction of heats of dilution are shown in the lower panels (as circles) and the solid lines represent the best least-square fits for the integrated data.

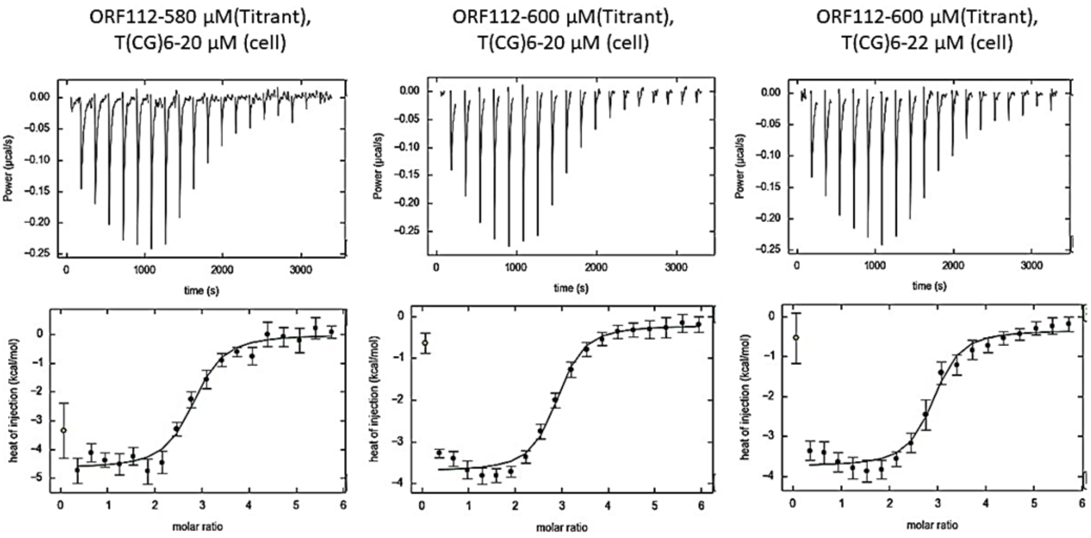


**Figure S8.** Isothermal titration calorimetry profiles for the titration of ORF112 R258A into a solution containing the $T(CG)_6$ oligonucleotide and $T(CG)_3$ oligonucleotide at various ratiometric concentrations of the protein and the oligonucleotide, respectively. The raw calorimetric titration data for the injections (2µl/injection) are shown in the upper panel and the integrated results after subtraction of heats of dilution are shown in the lower panels (as circles) and the solid lines represent the best least-square fits for the integrated data

**Table S1.** Table of thermodynamic parameters extracted from the fits.

| Prot | Conc (μM) | | N | K (×10[6] [1] and ×10[5] [2]) ($M^{-1}$) | ΔG | ΔH | TΔS |
|---|---|---|---|---|---|---|---|
| **ADAR1** | **P:1160 T(CG)$_3$: 72** | **1** | **1.07 ± 0.06** | **1.7 ± 0.95** | **-8.48** | **-2.13** | **-6.351** |
| | | **2** | **0.16 ± 0.04** | **2.1 ± 1.6** | **-7.26** | **-8.00** | **0.738** |
| | **P: 675 T(CG)$_3$:45** | **1** | **1.01 ± 0.3** | **1.1 ± 0.54** | **-8.23** | **-1.39** | **-6.841** |
| | | **2** | **0.27±0.3** | **4.0 ± 6.3** | **-7.64** | **-8.00** | **0.359** |
| | **P:1200 T(CG)$_3$: 70** | **1** | **0.992 ±0.3** | **1.2± 0.27** | **-8.29** | **-1.24** | **-7.05** |
| | | **2** | **0.282 ±0.3** | **5.7 ± 5.7** | **-7.85** | **-8.00** | **0.1506** |
| **ORF112** | **P: 675 T(CG)$_3$:45** | **1** | **1.16 ±0.03** | **3.9 ± 1.7** | **-8.98** | **-4.87** | **-4.113** |
| | | **2** | **0.356 ±0.08** | **0.13 ± 0.067** | **-5.61** | **-8.00** | **2.391** |
| | **P:814 T(CG)$_3$:43** | **1** | **1.54 ±0.09** | **1.5 ± 1.1** | **-8.43** | **-4.06** | **-4.36** |
| | | **2** | **0.319 ±0.3** | **0.082 ±0.15** | **-5.34** | **-8.00** | **2.66** |
| **ADAR1** | **P: 1160 T(CG)$_6$** | **1** | **1.76±0.06** | **5.7665±4.0804** | **-9.22** | **-0.125** | **-9.095** |
| | | **2** | **1.10±0.20** | **1.1450±0.29426** | **-6.9** | **-4.039** | **-2.859** |

**Appendix**

**Model 1:1 no transition B-to-Z**

Simple 1:1

Equilibrium constants:

Ka(A)=4.00e+06

Rate constants (1-forward, 2-reverse):

k1(A)=2.00e+04

k2(A)=5.00e-03

Chemical shifts:

w0(R)=0.0 /s (0.0 Hz)

w0(RL)=300.0 /s (47.8 Hz)

Base relaxation rates:

R2(R)=10.0 /s

R2(RL)=10.0 /s

Enthalpy difference from the base state:

dH(R)=0.0

dH(RL)=-200000.0

Total R concentration (*1000):

5.00 5.00 5.00 5.00 5.00 5.00 5.00 5.00 5.00 5.00 5.00 5.00 5.00 5.00 5.00 5.00 5.00 5.00 5.00 5.00 5.00 5.00 5.00

Ratio of total L to total R:

0.00 0.05 0.10 0.20 0.30 0.40 0.50 0.60 0.70 0.80 0.90 1.00 1.10 1.20 1.30 1.40 1.50 1.60 1.70 1.80 1.90 2.00 2.10

**Model 1:1 with B-to-Z assuming that Z-DNA is dominant (should recapitulate Model 1:1 with no transition)**

Simple 1:1 plus isomerisation

Equilibrium constants:

Ka(A)=4.00e+06

Ka(B)=1.00e-02

Rate constants (1-forward, 2-reverse):

k1(A)=2.00e+04

k2(A)=5.00e-03

k1(B)=1.00e-02

k2(B)=1.00e+00

Chemical shifts:

w0(R)=150.0 /s (23.9 Hz)

w0(R*)=0.0 /s (0.0 Hz)

w0(RL)=300.0 /s (47.8 Hz)

Base relaxation rates:

R2(R)=10.0 /s

R2(R*)=10.0 /s

R2(RL)=10.0 /s

Enthalpy difference from the base state:

dH(R)=0.0

dH(R*)=-70000.0

dH(RL)=-200000.0

Total R concentration (*1000):

5.00 5.00 5.00 5.00 5.00 5.00 5.00 5.00 5.00 5.00 5.00 5.00 5.00 5.00 5.00 5.00 5.00 5.00 5.00 5.00 5.00 5.00 5.00

Ratio of total L to total R:

0.00 0.05 0.10 0.20 0.30 0.40 0.50 0.60 0.70 0.80 0.90 1.00 1.10 1.20 1.30 1.40 1.50 1.60 1.70 1.80 1.90 2.00 2.10

**Model 1:1 with B-to-Z assuming that B-DNA is dominant**

Simple 1:1 plus isomerisation

Equilibrium constants:

Ka(A)=4.00e+06

Ka(B)=1.00e+02

Rate constants (1-forward, 2-reverse):

k1(A)=2.00e+04

k2(A)=5.00e-03

k1(B)=1.00e-02

k2(B)=1.00e-04

Chemical shifts:

w0(R)=150.0 /s (23.9 Hz)

w0(R*)=0.0 /s (0.0 Hz)

w0(RL)=300.0 /s (47.8 Hz)

Base relaxation rates:

R2(R)=10.0 /s

R2(R*)=10.0 /s

R2(RL)=10.0 /s

Enthalpy difference from the base state:

dH(R)=0.0

dH(R*)=-70000.0

dH(RL)=-200000.0

Total R concentration (*1000):

5.00 5.00 5.00 5.00 5.00 5.00 5.00 5.00 5.00 5.00 5.00 5.00 5.00 5.00 5.00 5.00 5.00 5.00
5.00 5.00 5.00 5.00 5.00

Ratio of total L to total R:

0.00 0.05 0.10 0.20 0.30 0.40 0.50 0.60 0.70 0.80 0.90 1.00 1.10 1.20 1.30 1.40 1.50 1.60
1.70 1.80 1.90 2.00 2.10
